# Leveraging *cis-* and *trans-*variants to improve protein expression level prediction for proteome-wide association studies

**DOI:** 10.64898/2026.05.28.728201

**Authors:** Rui Dong, Derek Lamb, Gao T. Wang, Andrew T. DeWan, Suzanne M. Leal

## Abstract

Since genetic effects are often mediated through proteins, the analysis of proteomic data can provide insights into disease etiology. However, most studies lack proteomic data. To address this problem, we developed TransCisPredict to perform proteome-wide association studies (PWAS) at a biobank scale. TransCisPredict reduces computational burden through linkage-disequilibrium block selection which facilitates incorporating *cis-* and *trans-*variants to predict protein expression and performs protein-phenotype association analyses. To account for differences in protein regulatory architecture, four prediction methods are used for weight estimation, i.e., BayesR, Elastic Net, LASSO, and SuSiE. Five-fold cross-validation (CV) is used to select the optimal method for each protein. Weight estimation was performed using White British UK Biobank study subjects (N=42,644) with proteomic and genotype array data. Of the 2,920 available protein expression levels, 2,339 could be predicted with a CV-R^2^>0.05 when *cis-* and *trans-*variants were used. Since most methods are limited to *cis-*variation, for comparison only *cis*-variants were used for prediction yielding 466 proteins with a CV-R^2^>0.05. A PWAS was performed for 2,339 predicted protein expression levels and type 2 diabetes (T2D) using White British UK Biobank study subjects without proteomic data (N=364,132) followed by two-sample Mendelian randomization using a method that controls for horizontal pleiotropy for validation. Forty proteins were associated with T2D and validated. For the 466 *cis-*only predicted protein expression levels, three proteins were associated with T2D and validated. Incorporating both *cis-* and *trans-*variation using TransCisPredict facilitates the prediction of many more proteins compared to using *cis-*only variants thereby increasing the power of PWAS.

## Introduction

Proteins play a central role in virtually all biological processes, acting as the primary functional mediators of genetic information(Sun et al. 2018; Anderson and Anderson 2002; Suhre et al. 2021). The study of how genetic variation influences RNA transcription or protein expression levels, often referred to as functional genomics, offers a critical lens through which to understand the molecular mechanisms underlying complex traits and diseases(Goh et al. 2018). While genome-wide association studies (GWAS) have uncovered thousands of genetic variants associated with complex diseases, the pathways through which these variants exert their effects remain unclear. Proteins, as downstream effectors of gene expression, provide a direct link between genotype and phenotype. However, most studies lack proteomic data making protein expression level prediction from genotypes crucial to study the protein-phenotype associations, making proteome-wide association studies (PWAS) an increasingly important research area(Suhre et al. 2021).

With advancements in high-throughput technology, large-scale studies have begun to generate proteomic data. The UK Biobank Pharma Proteomics Project (UKB-PPP) represents one of the most extensive efforts to date, with 2,923 measured plasma protein levels for >50,000 participants who also have genomic data. These data provide valuable insight into how genetic variation affects protein expression levels and the relationship between protein levels and disease etiology(Sun et al. 2023).

The generation of proteomic data is often limited by cost, sample availability, and technical constraints. Predicting protein levels from genomic data offers a solution(Wingo et al. 2020; Gedik et al. 2023). By leveraging genetic variants known to be associated with protein expression levels, these data can be predicted. This approach expands the application of proteomic data analysis to larger datasets, increasing statistical power to detect associations and improving the resolution to uncover novel signals.

Predicting protein levels from genetic data shares technical and conceptual similarities with prediction of transcriptomic data, where RNA expression levels are predicted based on genetic variation(Gedik et al. 2023). Although proteins can be the biological link between genetics and disease, similar to RNA expression levels, they are also impacted by environmental factors. Further, additional layers of genetically-driven regulation, such as post-translational modifications and degradation pathways, complicate predictions of protein levels(Brandes et al. 2020).

For RNA expression prediction, a reference cohort with both genomic and transcriptomic data are used to predict genetically driven RNA expression levels for a target cohort with genomic data. First, a model is developed to predict expression levels from the genomic data in the reference cohort. This model identifies a set of prediction weights between genomic variants and RNA expression levels while accounting for linkage disequilibrium (LD) between loci. Weights from this model are then used to predict RNA expression levels in a target cohort, followed by a transcriptome-wide association study (TWAS) to test for associations between predicted RNA expression levels and the phenotype of interest(Dai et al. 2023; Xue et al. 2020). Similarly, for protein expression levels, PWAS can be performed. A reference cohort with both genetic and protein expression level data are used to predict genetically driven protein levels. The estimated weights are then used in a target cohort with genomic data to predict protein expression levels that can be followed by a protein-phenotype association analysis. Accurate prediction of protein expression levels plays an important role in increasing the power of PWAS.

Numerous tools have been developed to predict RNA expression levels which include: PrediXcan(Gamazon et al. 2015), that utilizes elastic net regularization (Elastic Net) with an equal mixture of least absolute shrinkage and selection operator (LASSO) and Ridge regularization; FUSION(Gusev et al. 2016), that uses a Bayesian sparse linear mixed model; and TIGAR(Nagpal et al. 2019), a nonparametric approach. Each of these methods imposes different assumptions about the distribution of quantitative trait loci (QTLs) in the prediction model. Various methods can also be combined into an omnibus approach, accommodating the different genetic architecture across RNAs(Dai et al. 2023). These tools have also been applied to proteomic data, leveraging the similarities between RNA and protein expression level data(Schubert et al. 2022; Hu et al. 2024a; Liu 2025).

A current limitation of predicting RNA and protein expression levels is that most methods only incorporate *cis-*variants, due to the computational burden and methodological limitations of also incorporating *trans-*variants(Powder 2020). By reducing the dimension of the predictor set used for transcript or protein prediction to *cis-*variants, computational efficiency is gained at the expense of loss of *trans* information. This approach is problematic, since for both RNA and protein expression levels, *trans*-variants account for the majority of heritability, i.e., for RNA expression levels the mean (μ) expression (e) QTL heritability is 6% for *cis* and 26% for total heritability(Ouwens et al. 2020) and for protein (p) QTL heritability is 7.5% for *cis* and 19.5% for total heritability(Sun et al. 2023).

Incorporating *trans*-variation in protein prediction models captures critical information on the genetic drivers of protein expression levels. *Cis-*variation within a gene may regulate protein expression levels through direct impact on the stability of the protein, or they may lie within the gene’s promoter or another regulatory element, e.g., enhancers and repressors. However, variants within transcription factors that bind these regulatory elements or regulate proteins post-translation(Chen et al. 2024b) are typically *trans* to the target gene. Also, given that *cis*-variants do not always modulate protein expression levels, and *cis-*and *trans*-variants can have inverse effects on protein levels(Goncalves et al. 2012; Landry et al. 2005; Metzger et al. 2017; Signor and Nuzhdin 2018), using both *cis-* and *trans*-variants in the prediction of protein expression levels is critical. Although some methods for transcriptomic data have been developed that incorporate trans-variants, e.g., BGW-TWAS(Luningham et al. 2020) and transTF-TWAS(He et al. 2025), they are limited in their applications. TransTF-TWAS only includes *trans*-variants linked to a transcription factor, disregarding other *trans* information, while BGW-TWAS imposes a single variable selection approach rather than more flexible methods that can accommodate a range of genetic architectures. While recent studies have begun to incorporate trans-pQTLs to improve protein prediction, they either use univariate statistical association test results which focus on marginal rather than joint effects(Mosley et al. 2018; Xu et al. 2023; Western et al. 2024; Yang et al. 2025a), or utilizes a non-linear XGBoost machine learning framework that lacks the direct interpretability and portability of additive effect sizes (Sigurdsson et al. 2025). There remains a need for a flexible framework that systematically optimizes prediction models across varying genetic architectures. Most current approaches rely on a single regularization method, a fixed machine learning architecture, or pick the “lead SNP” within a window, which may not be optimal for the diverse regulatory landscapes of different proteins. Our approach, TransCisPredict, builds upon these approaches by evaluating a range of flexible variable selection models, to account for proteins that may range from having sparse, large-effect *cis* regulation to those with highly polygenic trans-acting components.

TransCisPredict incorporates *cis-* and *trans*-variants by selects a subset of LD blocks to decrease computational burden and also incorporates four flexible variable selection methods, Bayesian mixture of normals (BayesR), Elastic Net(Zou and Hastie 2005), LASSO(Tibshirani 1996), and sum of single effects regression (SuSiE)(Wang et al. 2020) to account for protein-specific genetic architectures. The optimal method for each protein is selected via cross validation (CV) to increase prediction accuracy (Figure 1). This ensures that the specific regulatory landscape of each protein—whether driven by a few large-effect trans-loci or many small-effect variants—is accurately captured. We applied TransCisPredict at biobank-scale, first demonstrating that within the UKB-PPP, the inclusion of *trans*-variants in the prediction models substantially improves protein prediction compared to a *cis*-only model allowing for the prediction of 5x the number of proteins.

**Figure 1.**
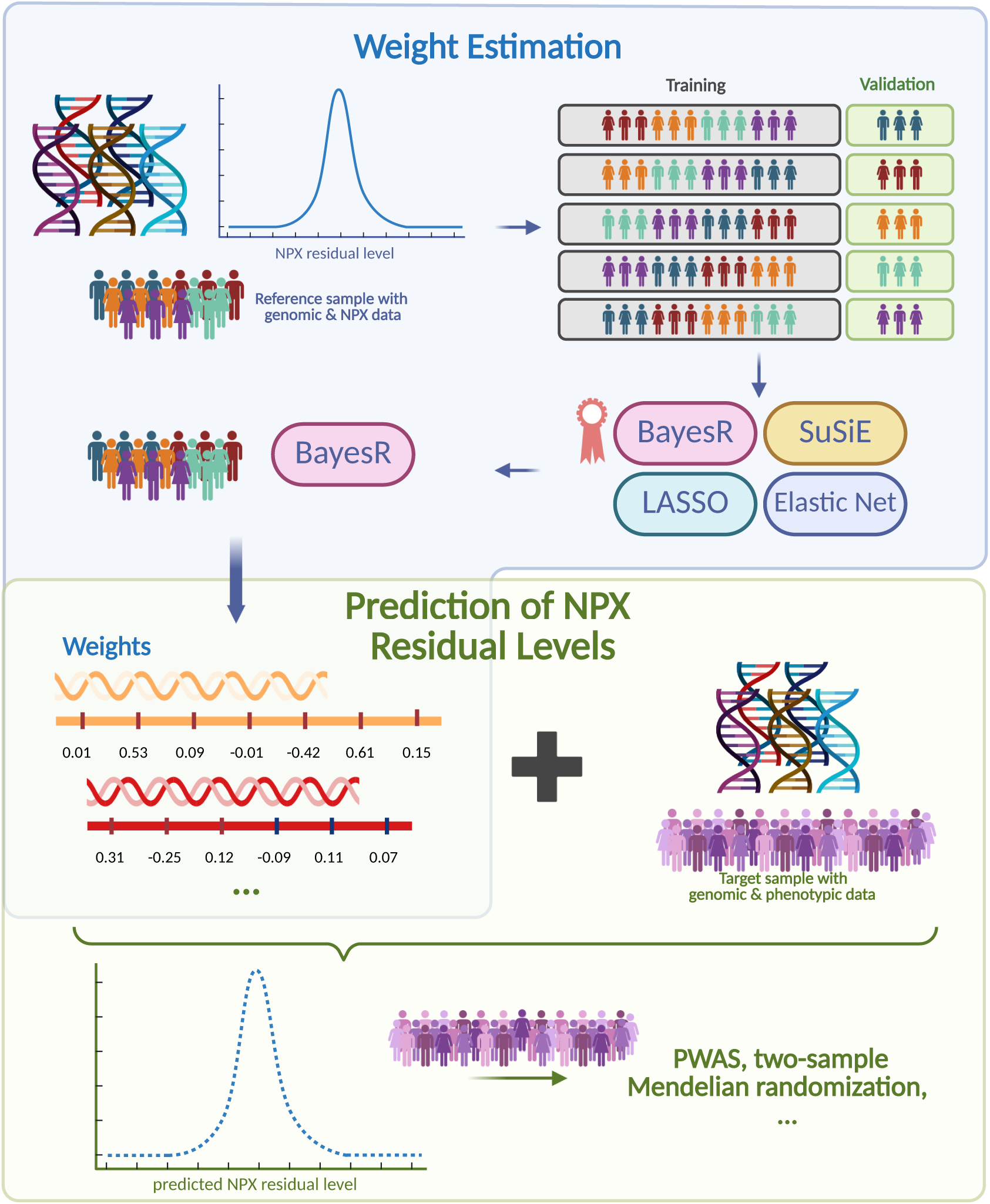
Overview of the TransCisPredict workflow. A reference sample with genomic and proteomic data is first used to select LD blocks for each protein which are used in weight estimation (see Figure 4 for additional details). Using all variants in the selected LD blocks, five-fold cross-validation is performed using four methods (BayesR, Elastic Net, LASSO, and SuSiE), and the method with the highest CV-R² is selected. The selected method is then applied to all reference samples to estimate weights for the protein. The weights are then applied to target samples with genomic data to generate predicted protein residual levels. Lastly, a protein-phenotype association analysis is performed using the predicted protein residual levels for a dichotomous trait (quantitative traits such as lipid levels can also be analyzed). This workflow would usually be performed across all proteins as shown in the volcano plot. The arrow points to the result for the example protein.

Using TranCisPredict, a PWAS was performed for type 2 diabetes (T2D) using White British UK Biobank study subjects without proteomic data. For the detected protein-T2D associations it was determined if the findings could be validated using two-sample Mendelian randomization (MR)-pleiotropy residual sum and outlier (PRESSO)(Verbanck et al. 2018) that controls for horizontal pleiotropy. For protein expression levels that were predicted using *cis-*only variants, only three proteins were associated with T2D and validated. In contrast, when prediction was performed using *cis-* and *trans-*variants 13.3X more proteins (N=40) were associated with T2D and validated. We also analyzed measured protein expression levels and performed a direct PWAS (dPWAS) for T2D. The PWAS had a higher percent of associated and validated proteins (48.2%) compared to the dPWAS (33.2%). Additionally, for the PWAS we identified five validated T2D associated proteins which were not associated in the dPWAS. For prediction of protein expression levels, inclusion of *cis-* and *trans-*variants not only increases the number of proteins which can be predicted but also facilitates new protein-phenotype associations.

## Results

### Number of cis-only and cis– and trans-LD blocks included in the prediction

For the *cis-*only analysis, the number of LD blocks included for the prediction of each NPX residual level ranged from 1 to 5 (μ=2.27). For the *cis-* and t*rans-*analysis, the number of LD blocks used for the prediction of NPX residual levels ranged from 274 to 653 (μ=459.09) (Table 1).

**Table 1.** Number of LD blocks used for NPX residual level prediction for *cis-*only and *cis-* and *trans-* analyses.

| # LD blocks |  | Minimum | 25% Quantile | Median | Mean | 75% Quantile | Maximum |
| --- | --- | --- | --- | --- | --- | --- | --- |
| *cis*-only |  | 1 | 2 | 2 | 2.27 | 3 | 5 |
| *cis*- and *trans*- | Total # LD blocks | 274 | 404 | 454 | 452.09 | 495 | 653 |
|  | # *cis*-LD blocks | 0 | 1 | 2 | 1.55 | 2 | 5 |
|  | # *trans*-LD blocks | 272 | 402 | 453 | 450.54 | 494 | 651 |

Abbreviations: #-number of; LD-linkage disequilibrium; NPX-normalized protein expression

### Weights Estimation

A total of 466 NPX residual levels (15.96% of all available proteins) using *cis-*only variants and 2,339 NPX residual levels using the *cis-* and *trans-*variants (80.10% of all available proteins) could be predicted with a CV-R^2^>0.05 (Tables 2 and S1). For the 466 NPX residual levels predicted using *cis-*only variants, the CV-R^2^ μ=0.181 (Table S2), and for the NPX residual levels predicted using *cis-* and *trans-*variants, the CV-R^2^ μ=0.125. For prediction using *cis*-only variants, SuSiE was most often selected to perform the NPX residual level prediction (45.92%), followed by BayesR (30.47%). For prediction performed using *cis-* and *trans-*variants, 93.12 % of the 2,339 proteins were predicted using BayesR, and 4.32%, 1.71%, and 0.86% for SuSiE, LASSO, and Elastic Net, respectively (Tables 3 and S1). For the weight estimation using *cis-* and *trans-*variants 60 protein had the highest CV-R^2^ when either LASSO or Elastic Net was used for weight estimation, the average decrease in CV-R^2^ is 0.047 (median: 0.032) if only BayesR and SuSiE were used, and 23 of these proteins would have had a CV-R^2^≤0.05 (Table S3).

**Table 2.**
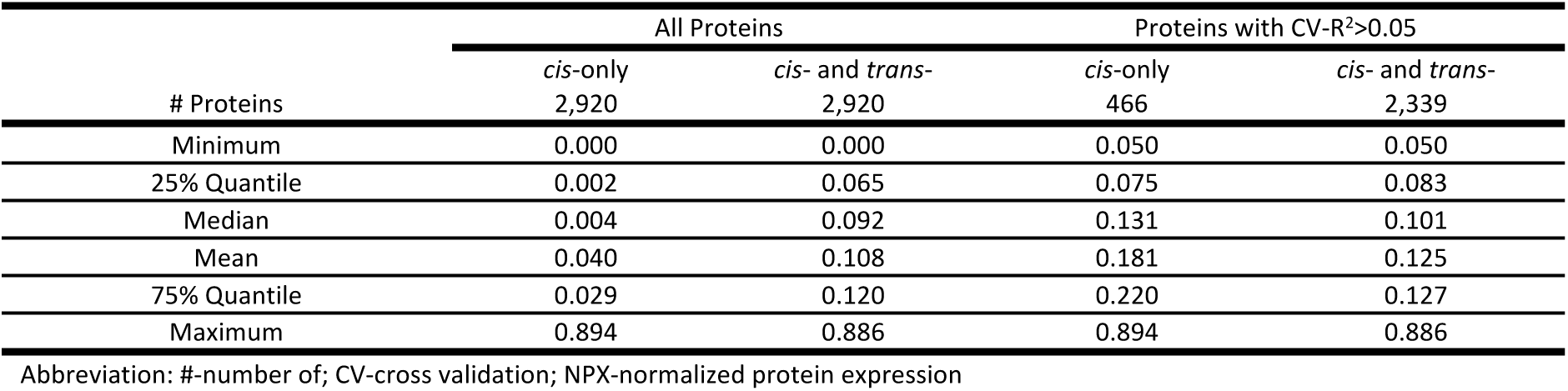
CV-R^2^ for predicted NPX residual levels.

**Table 3.** Optimal method selected to predict NPX residual levels (CV-R^2^>0.05)

| Selected Method | # Proteins | BayesR | SuSiE | LASSO | Elastic Net |
| --- | --- | --- | --- | --- | --- |
| *cis*-only | 466 | 142 (30.47%) | 214 (45.92%) | 49 (10.52%) | 61 (13.09%) |
| *cis*- and *trans*- | 2,339 | 2,178 (93.12%) | 101 (4.32%) | 40 (1.71%) | 20 (0.86%) |

Abbreviations: #-number of; BayesR-Bayesian mixture of normals; CV-cross validation; LASSO-least absolute shrinkage and selection operator regularization; NPX-normalized protein expression; SuSiE-sum of single effects regression

The evaluation of the threshold for the FDR q-value to select *cis-* and *trans-*LD blocks for estimating prediction weights demonstrated that an FDR q-value of <0.2 led to the ability to predict more NPX residual levels than using an FDR q-value <0.1, i.e., N=2,339 compared to N=1,826 predicted NPX residual levels, an increase of 28.09% (Tables S1 and S4).

### Association Analysis of Predicted and Measured NPX Residual Levels and Type 2 Diabetes

The analysis of 2,339 predicted NPX residual levels for the N=19,964 T2D cases, and N=280,453 controls yielded a significant association (p-value<1.71×10^−5^) for 83 predicted NPX residual levels (Figure 2, Tables 4, S5, and S6). When the same dataset was analyzed but using the 466 NPX residual levels predicted using *cis-*only variants, three predicted NPX residual levels were significantly associated with T2D (Tables S7 and S8).

**Figure 2.**
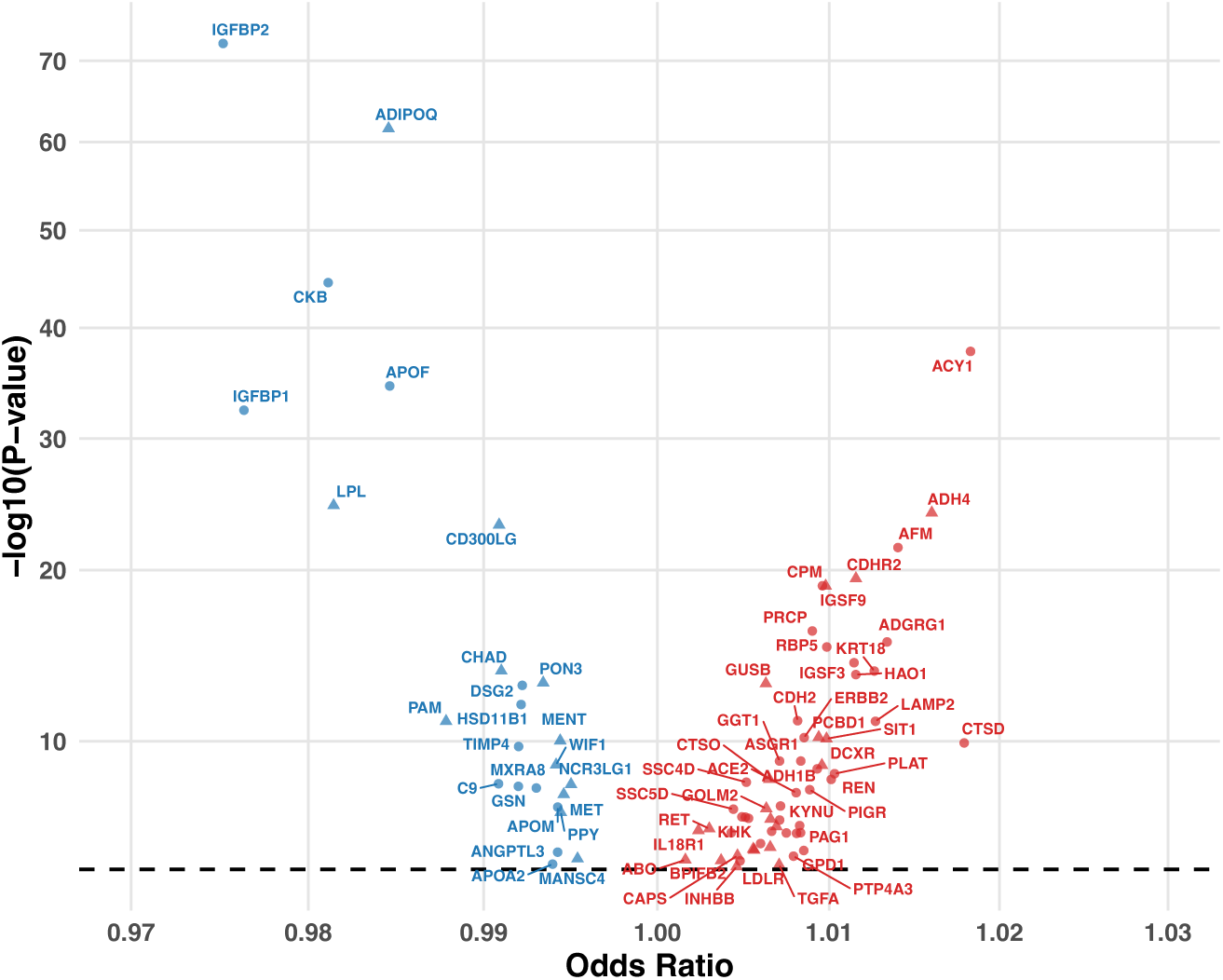
Proteome-wide association study for type 2 diabetes. Volcano plot of 83 proteins significantly associated with T2D using predicted NPX residual levels. The dashed line indicates the Bonferroni-corrected significance level (p-value=1.71 × 10⁻⁵). Triangles denote proteins that were associated with T2D and validated using-MR-PRESSO (p-value<0.05); circles denote proteins that were associated with T2D but could not be validated. Red circles and triangles indicate an increase and blue circles and triangles indicate a decrease in NPX residual levels associated with T2D risk.

**Table 4.** Type 2 Diabetes PWAS and dPWAS results and validation.

| NPX Residual Source | # Significant proteins<sup>a</sup> | # Validated<sup>b</sup> | Proportion | 95% CI |
| --- | --- | --- | --- | --- |
| Prediction *cis*- and *trans*- variants | 83 | 40 | 0.482 | 0.374 – 0.589 |
| Unique for predicted NPX residual levels | (8) | (5) | (0.625) | (0.290 – 0.961) |
| Prediction *cis*-only variants | 3 | 3 | 1 | - |
| Unique for predicted NPX residual levels | (3) | (3) | (1) | (-) |
| Measured | 1,052 | 380 | 0.361 | 0.332 – 0.390 |
| Among 2339 predictable proteins | (854) | (322) | (0.377) | (0.345 – 0.410) |
| Among 581 unpredictable proteins | (198) | (58) | (0.293) | (0.230 – 0.356) |

<sup>a</sup>Significant in PWAS or dPWAS (p-value<1.71×10<sup>-5</sup>). <sup>b</sup>Validated using MR-PRESSO (p-value<0.05). Abbreviations: #-number of; CI-confidence interval; dPWAS-direct proteome-wide association studies; MR-PRESSO-Mendelian randomization pleiotropy residual sum and outlier; NPX-normalized protein expression; PWAS-proteome-wide association studies

For the 2,920 measured NPX residual levels, we analyzed N=2,556 T2D cases and 32,094 controls and detected 1,052 significantly associated NPX residual levels (Tables 4, S9, and S10). For these significantly associated NPX residual levels, 854 could be predicted using *cis-* and *trans-*variation (Tables S9 and S10), but 198 of these proteins could not be predicted (Tables S11 and S12).

For the 83 predicted NPX residual levels that were significantly associated with T2D, eight predicted NPX residual levels: ABO, APOA2, CAPS, DSG2, LDLR, MANSC4, PAM, and SFRP4, were unique to analyses using predicted NPX residual levels and were not detected when measured NPX residual levels were analyzed. Three of the proteins, ABO, MANSC4, and PAM, were also detected to be associated with T2D when protein prediction was performed using *cis-*only variants.

### Two-Sample Mendelian Randomization

Of the 83 predicted NPX residual levels that were associated with T2D, 40 were validated using MR-PRESSO [0.482, 95% confidence interval (CI): 0.374-0.589] (Figure 3, Tables 4 and S5). All three proteins which were predicted using *cis-*only variants and associated with T2D were validated using MR-PRESSO (Table S7). For the 1,052 measured NPX residual levels that were associated with T2D, 380 (0.332, 95% CI: 0.332-0.390) of them were validated using MR-PRESSO (Tables S9 and S10). For the eight proteins for which an association with T2D was only observed for predicted NPX residual levels, five (0.625, 95% CI: 0.290-0.961) were validated using two-sample MR-PRESSO (ABO, CAPS, MANSC4, PAM, and SFRP4).

**Figure 3.**
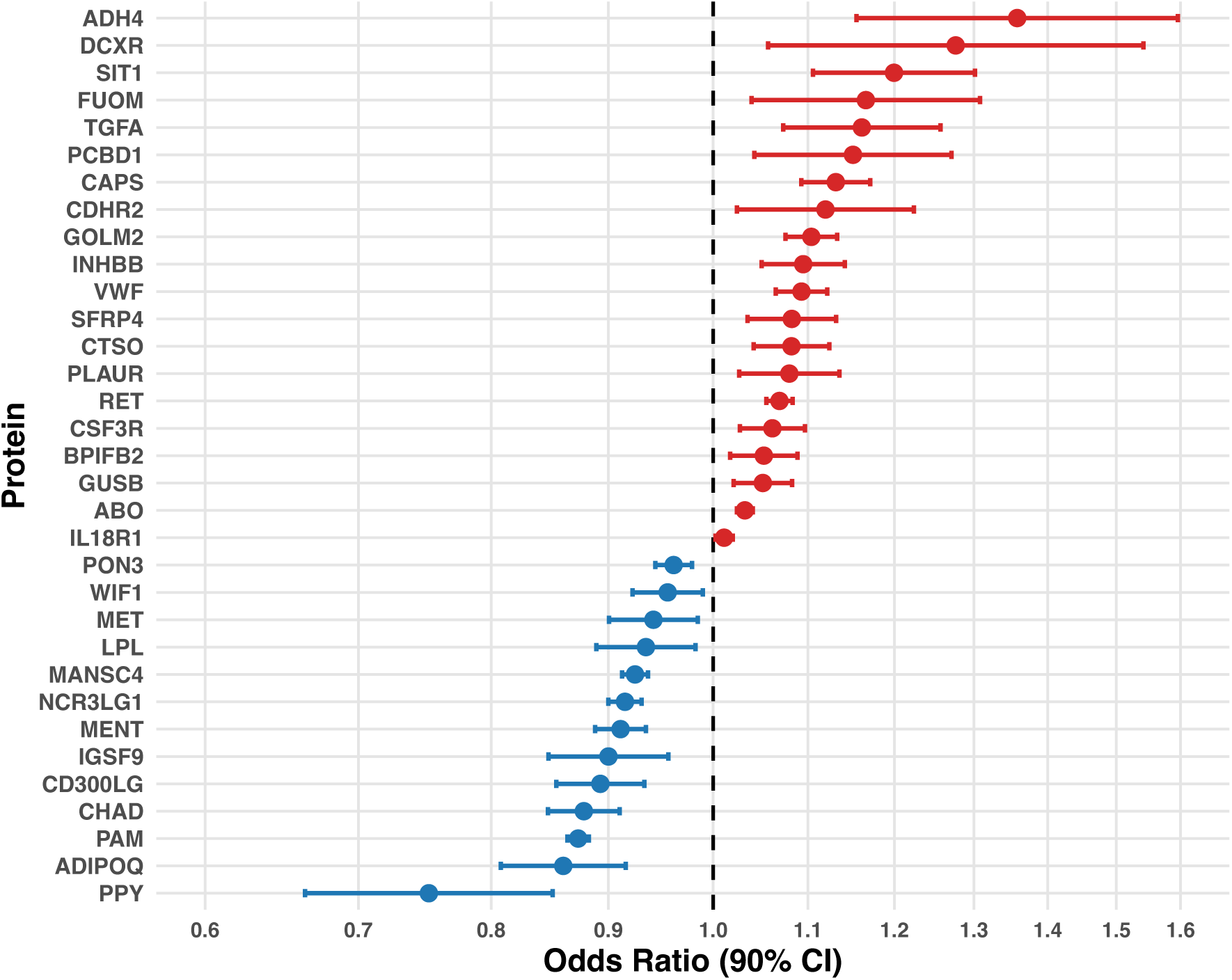
Validated protein-type 2 diabetes associations using Mendelian randomization. The 40 proteins which were validated using MR-PRESSO (p-value<0.05) with effect sizes and 95% confidence intervals.

## Discussion

We developed TransCisPredict, a framework to perform PWAS, that incorporates both *cis-* and *trans-*variants to perform protein expression level prediction and protein-phenotype association analysis at biobank-scale. Compared to *cis*-only models, including *trans-*variants substantially improved protein expression level prediction and greatly increased the number of predictable proteins. Here we could predict 5.0x the number of protein expression levels using *cis-* and *trans-*variation compared to limiting prediction to *cis-*only variants. The ability to predict more protein expression levels impacted the T2D association results, i.e., we were able to detect associations and validate the findings using two-sample MR for 13.3x as many protein expression levels when prediction was performed using *cis-* and *trans-*variation compared to performing prediction using *cis-*only variants. Our findings align with recent reports suggesting that the inclusion of trans-acting variants significantly increases the heritability explained for the plasma proteome(Chen et al. 2024a). However, TransCisPredict offers unique computational advantages. By partitioning the genome into independent LD blocks and applying a relaxed FDR threshold (q < 0.2) for block selection, we capture a larger proportion of trans-heritability compared to methods that only include genome-wide significant SNPs. Additionally, BayesR outperformed other methods for 93.1% of proteins for the analysis which includes *cis-* and *trans*-variants highlights the complex, mixture-of-normals architecture of trans-protein regulation, a nuance not fully explored in previous PWAS methodologies.

TransCisPredict represents a methodological advance for protein expression level prediction. Rather than first identifying a stringent set of significant *trans-*variants and then using them for prediction, we partitioned the genome into 1,703 independent LD blocks and selected regions to be used in weight estimation. This strategy focused on maximizing prediction accuracy while reducing computational burden. The approximate independence between LD blocks allows for the variable selection methods, e.g. BayesR to analyze hundreds or thousands of variants at a time, instead of all variants on a chromosome, drastically reducing the time and memory required for computation. To further reduce computational burden, we selected a subset of LD blocks to include in the analysis by using a relaxed inclusion criterion (FDR q-value<0.2), which led to a ∼75% reduction in the number of LD blocks that were analyzed for weight estimation compared to analyzing all LD blocks.

This study used genotype array data for computational efficiency, which does not capture all genetic variation present in the genome. The 646,469 array variants that passed quality control provide substantial coverage of common variation through tagging of variants that were not genotyped. Other genetic data could be used, such as imputed variants or whole-genome sequence data. These datasets would include a larger number of variant sites, including rare variants, which have been shown to impact protein expression levels(Dhindsa et al. 2023). These variants, including rare variants, can be assigned to LD blocks, and association testing can then be performed to determine which LD blocks should be included in weight estimation. However, estimating weights using denser variant data comes with substantially higher computational memory and time costs.

The CV framework, incorporated in TransCisPredict, selects the optimal method for each protein, preventing overfitting and ensuring predictions generalize well to other datasets—a crucial consideration given the often-unknown regulatory landscape governing protein expression levels—making this framework well-suited to predict a variety of proteins. We employed a cross-validated CV-R² threshold of 0.05 for model inclusion. This threshold is consistent with, or more stringent than, established transcriptomic and proteomic prediction studies [Gusev et al. 2016, Wingo et al. 2020], ensuring that the models capture a robust genetic signal. While an R² of 0.05 represents a minority of the total protein variance, in the framework of PWAS, these models serve as valid genetic instruments that are inherently shielded from the environmental confounding and reverse causation often present in dPWAS. TransCisPredict is flexible and is not limited to BayesR, Elastic Net, LASSO, and SuSiE; fewer or more variable selection methods can be used and as new methods are developed, they can also be incorporated. BayesR for most proteins outperforming the other three methods in *cis-* and *trans*-variant penalized regression models—provides insight into the regulatory landscape of the human proteome. Our results show that BayesR increases the mean prediction R² by 75.7% compared to the best of the other three methods for the *cis*– and *trans*-variant analysis. This improvement suggests that protein expression is governed by an architecture where a few high-effect cis-acting variants coexist with a moderate number of *trans*-acting variants with varying effect sizes. This supports the ‘omnigenic’ perspective of protein regulation, where distal genetic variation contributes significantly to phenotypic variance through interconnected molecular networks.

Five validated associations were observed for predicted NPX residual levels and T2D, i.e., ABO, CAPS, MANSC4, PAM, and SFRP4, but not detected in the dPWAS. The ability to detect associations with predicted protein expression levels and not those that are measured may occur because there is a strong genetic component that is modulating protein expression levels and the sample size using predicted NPX residual levels was >8.5x larger than for the measured NPX residual levels, increasing the power to detect associations. These five proteins display evidence of being involved in T2D disease etiology. The association between increased ABO protein expression levels and T2D risk is consistent with prior evidence that non-O blood groups exhibit higher von Willebrand factor and inflammatory markers, which promote insulin resistance and increase diabetes susceptibility(Meo et al. 2016; Legese et al. 2020; Yuan et al. 2023). The positive association between calcyphosine (CAPS) and T2D risk aligns with recent single-cell studies identifying CAPS as a marker of β-cell dedifferentiation, where it is re-expressed as mature β-cells revert to an immature state with impaired insulin secretion(Wei et al. 2025). Higher MANSC4 protein expression levels are associated with reduced T2D risk, potentially through its predicted function as a serine-type endopeptidase inhibitor(Yuan et al. 2023; Loesch et al. 2025; Yang et al. 2025b). Reduced PAM protein expression or activity levels are associated with decreased insulin content, and impaired insulin secretion from pancreatic β-cells contributes to increased T2D risk(Thomsen et al. 2018; Giontella et al. 2025). Elevated SFRP4 protein expression levels are associated with increased T2D risk, potentially through reduced insulin secretion as SFRP4 has been shown to decrease β-cell Ca²⁺ channel expression and suppress insulin exocytosis(Dai et al. 2023; Anand et al. 2016; Sánchez-Pozos et al. 2024). While the total number of new associations may appear modest, these represent ‘genetically-driven’ associations which are high-priority candidates for therapeutic intervention. Furthermore, the utility of TransCisPredict is most pronounced in scenarios where proteomic data is scarce or unavailable. As GWAS meta-analyses reach sample sizes in the millions, the power to leverage even modest genetic prediction (e.g., CV-R²) to discover novel disease biology will become increasingly vital.

Our application of TransCisPredict to T2D GWAS data demonstrated a clear trade-off between model specificity and sensitivity. Cis-only models exhibited a perfect validation rate (100%) in MR analysis, likely due to the large effect sizes and direct biological mechanisms inherent to cis-regulation. However, the inclusion of trans-LD blocks expanded the number of validated protein-trait associations from 3 to 40. This suggests that while trans-signals may be more susceptible to environmental noise or horizontal pleiotropy—resulting in a lower percentage of validation (48%)—their inclusion is critical for capturing the full spectrum of protein-to-disease associations. For the majority of the plasma proteome, the ‘missing heritability’ lies in the *trans*-components; therefore, incorporating these variants is essential for maximizing the discovery power of PWAS frameworks.

While *cis-*pQTLs are often preferred for causal inference due to a reduced risk of horizontal pleiotropy, our framework leverages MR-PRESSO to rigorously filter for pleiotropic outliers, thereby enabling the inclusion of informative *trans*-variants that would otherwise be discarded. Although MR-PRESSO was originally developed for complex traits where the instrumental variable includes multiple loci, its application here serves as a stringent secondary validation layer to complement our proteome-wide screening. By identifying 40 validated protein-T2D associations compared to only three using *cis*-only models, we demonstrate that a cautious integration of *trans*-acting variation is essential for capturing the full regulatory landscape of the proteome.

This study has limitations. The analysis only included White British due to data availability in the UK Biobank-PPP, since for this dataset there are very limited numbers of individuals of non-White European ancestry, e.g., South Asians. It is not clear how analysis of non-European ancestry groups impacts the ability to predict protein expression levels and if weights generated from one population are transferable across ancestry groups.

Testing for associations with protein expression levels, rather than relying solely on results from GWAS, offers critical advantages in identifying potential therapeutic interventions. While GWAS identifies genetic variants associated with diseases, protein-level associations provide a direct link to the functional mediators of biological processes. Most importantly, proteins represent the primary targets of therapeutic agents, making protein-level associations particularly valuable for drug discovery(Geddes-McAlister et al. 2022; Ghoussaini et al. 2023). By identifying proteins with genetically driven levels in plasma that are correlated with disease phenotypes, PWAS and dPWAS pinpoints molecules that can be directly targeted, potentially guiding the development of novel compounds or the repurposing of existing therapeutics.

The UK Biobank currently has the largest sample of proteomic data for an individual study and will soon have measured proteomic data for almost all 500,000 study subjects. Given the sample size of the UK Biobank and availability of phenotypic data, the measured protein expression levels provided excellent power to detect associations. Many studies have genomic data, but do not have proteomic data or only have proteomic data available for a subset of study subjects. For example, All of Us(All of Us Research Program Investigators et al. 2019), a population-based United States study, currently has genotype array data for 447,300 and whole-genome sequence data for 414,840 study subjects, but it plans to only release proteomic data for 10,043 study subjects. Even for well powered dPWAS, it can still be advantageous to also perform a PWAS since the genetic component of protein expression level is being predicted which can sometimes allow for the detection of additional protein-phenotype associations which can be validated as demonstrated in our analysis of T2D. To perform PWAS, TransCisPredict offers a flexible framework to perform analyses on a biobank-scale.

## Material and Methods

### Data Access and Ethical Approval

This research was conducted using UK Biobank data (application number 32285) and their Research Analysis Platform. The UK Biobank study was conducted under generic approval from the National Health Services’ National Research Ethics Service. The present analyses were approved by the Institutional Review Boards at Yale University (2000026836) and Columbia University (AAAS3494).

### Quality Control of Genotype-Array Data and Samples

The UK BiLEVE array and the UK Biobank Axiom array(Bycroft et al. 2018) were used to assay samples from UK Biobank. The intersection of these arrays contains 733,322 autosomal variants. Genotype quality control was performed for White British study subjects (f.22006) (N=409,460) after removing participants that: 1) are outliers for genotype heterozygosity or missing genotype data (f.22027) (N=727); 2) have inconsistent self-reported sex (f.31) and genomic sex (f.22001) (N=307); 3) have a sex chromosome aneuploidy (f.22019) (N=407); or 4) withdrew their consent to participate in the study (N=102). Variant sites were removed that had excess missing genotypes, i.e., variants with a minor allele frequency (MAF) ≥0.05 with a call rate <0.95 and variants with a MAF <0.05 with a call rate <0.99. Variants were then tested for deviations from Hardy-Weinberg equilibrium (HWE) among unrelated study subjects. To obtain unrelated study subjects, KING(Manichaikul et al. 2010) was used to estimate kinship coefficients, and for related pairs (kinship coefficient>0.044), only one individual was retained. Those variant sites that deviated from HWE (p-value<1.0×10^−15^) were removed, leaving a total of 646,469 variants available for analysis.

### Protein Expression Level Data

The UKB-PPP project generated plasma protein expression levels for 53,018 UK Biobank participants utilizing the Olink Explore 3072 platform to measure 2,923 proteins and reported normalized protein expression (NPX) levels (f.30900). Three proteins, GLIPR1, NPM1, and PCOLCE, were missing >70% of their observations and were removed from further analysis. Of the 53,018 study participants with proteomic data, 43,672 are of White British ancestry and passed quality control. Residuals were generated for the NPX levels by regressing out age (f.21003), sex (f.31), age × sex, body mass index (BMI) (f.21001), and the first 20 genetic principal components (PCs). PCs were generated using FlashPCA***(Abraham and Inouye 2014)*** using 226,844 variants that were pruned for LD (r^2^<0.1). Of these study subjects, 42,644 are unrelated (kinship coefficient<0.044) and were used for weight estimation.

### Weight Estimation

Weight estimation to predict NPX residual levels was performed using two strategies: 1) using *cis*-only variants; and 2) *cis-* and *trans*-variants.

To select the *cis*– and *trans-*variants to include in the weight estimation, the 2,920 NPX residual levels were first tested for genetic associations using the unrelated White British study subjects using linear regression. Each NPX residual level was tested for an association with each of the 646,469 variants that lie on 1,703 LD autosomal blocks as defined for Europeans(Berisa and Pickrell 2016). Covariates included in the analysis were age, sex, age x sex, BMI, and 20 genetic PCs.

For the *cis-* and *trans-*variant analysis, a Benjamini-Hochberg false discovery rate (FDR) was used to select LD blocks to be used in weight estimation while accounting for multiple testing within each LD block (Figure 4). Two different q-values <0.1 and <0.2 were evaluated. A q-value <0.2 was selected because it allowed for the better prediction of protein expression levels and was used for all subsequent analyses (see Tables S1 and S4). For the *cis-*only analysis, LD blocks that span the gene coding the protein ±1MB (Figure 5) were used to estimate weights(Sun et al. 2023). Once a block was selected, all genetic variants located within that block’s boundaries were included as features for model training. We deliberately avoided restricting the input to ‘lead SNPs’ to prevent the loss of secondary signals or poorly tagged causal variants.

**Figure 4.**
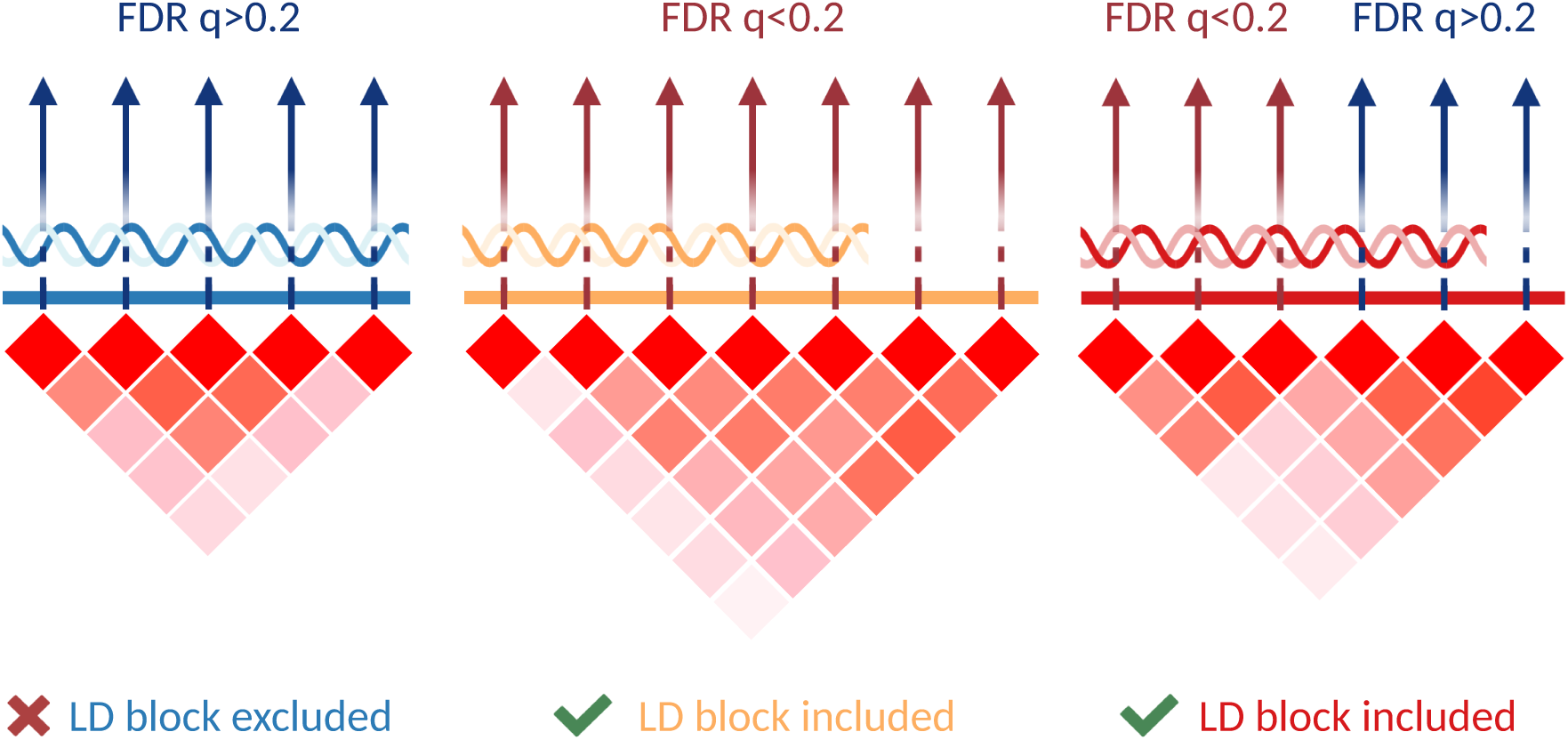
Selection of LD blocks based on FDR q-values. LD blocks are included in subsequent analysis if at least one variant has FDR q<0.2. The heatmaps show LD (r²) among variants. Blue arrows indicate variants with an FDR q>0.2, red arrows indicate an FDR q<0.2. Left panel: LD block excluded (all variants FDR q>0.2) from weight estimation. Middle and right panels: LD blocks included (≥1 variant with FDR q<0.2) in weight estimation.

**Figure 5.**
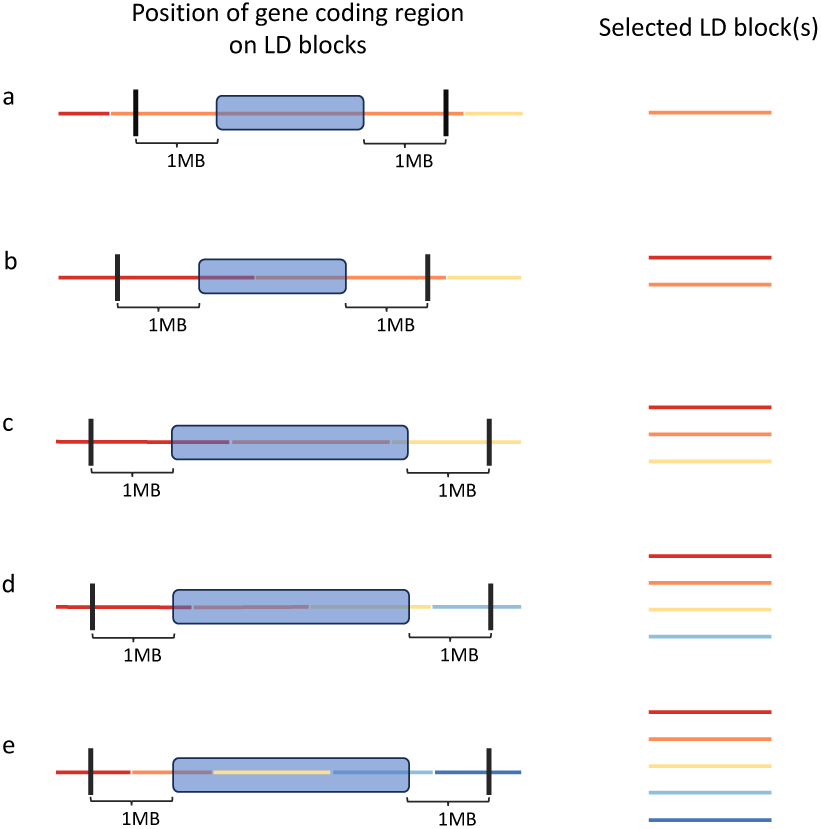
Definition of cis-regulatory LD blocks. Any LD block (red, orange, yellow, sky blue, and dark blue lines) overlapping with the gene coding region (blue box) ± 1Mb is considered a *cis-*regulatory LD block. Five scenarios (a-e) illustrate different configurations based on the position of the gene coding region relative to LD block boundaries.

TransCisPredict employs four variable selection methods to accommodate different genetic architectures: BayesR, Elastic Net, LASSO, and SuSiE. LASSO and SuSiE both assume sparse signals, with LASSO using an L1 penalty that drives most coefficients to zero and SuSiE modeling a fixed number of maximum independent causal effects. Bayes R and Elastic Net are more data-adaptive, compared to LASSO and SuSiE. BayesR uses a mixture distribution that estimates the proportions of null, small, medium, and large effects directly from the data, providing flexibility to model genetic architectures spanning sparse to highly polygenic traits. For Elastic Net, the regularization strength is tuned by CV but the mixing parameter (alpha=0.5) is fixed. We relied on the internal regularization and variable selection mechanisms of the four evaluated models (BayesR, SuSiE, LASSO, and Elastic Net) to characterize the local genetic architecture for each selected LD block. This approach ensures that the models can distinguish between sparse, single-variant effects and more complex, polygenic configurations within a given trans-locus.

Five-fold CV was applied to select the optimal method for each protein (Figure 1). The sample of 42,644 unrelated White British study subjects were divided into five groups without replacement for each of the 2,920 proteins, and four of the groups (80% of the sample) were used to estimate weights, these weights were then used to predict NPX residual levels in the fifth group (20% of the sample) and the CV-R² was estimated. This procedure was repeated, each time using a different one of the five groups to perform NPX residual level prediction and estimate the CV-R^2^ with the remaining 80% of samples used to estimate weights. For each method, we computed the Pearson correlation between the measured and predicted NPX residual levels, i.e., CV-r, and the proportion of variance in the measured NPX residuals explained by predicted values, i.e., CV-R². For each protein, the method with the highest average CV-R² across the five groups was selected(Gusev et al. 2016; Hu et al. 2024b) and this optimal method was then applied to the entire sample (White British N=42,644) to estimate the final weights. For each protein both *cis-*only and *cis-* and *trans-* analyses were performed. For the CV procedure the study subjects to be included in each of the five groups were randomly selected. Only proteins with a CV-R^2^ >0.05 and CV-r>0 (hereafter denoted as CV-R^2^ >0.05) were included in subsequent analyses.

### Prediction of Protein Expression Levels

NPX residual levels were predicted for 364,132 White British study subjects (related and unrelated) without measured NPX levels using the final weights by summing up the product of the genotype variants and the corresponding weights. For each protein, the weights for each genetic variant are denoted as ***w*** = (*w*_1_, …, *w_M_*) where M is the number of genetic variants. For the *i*-th individual (*i* = 1,2, …, *N*), we denote the genotype values as *G*_i⋅_ = (*G*_i1_, …, *G_iM_*) where *G_ij_* is the number of effect alleles that the *i*-th individual carries for the *j*-th variant. The predicted NPX residual level for the *i*-th individual is:

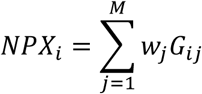

The above prediction was performed twice, once for *cis-* and *trans-*variation and then for *cis-*only variation.

### Association Analysis of Predicted and Measured NPX Residual Levels and Type 2 Diabetes

Using the predicted NPX residual levels based on prediction performed using *cis-* and *trans-*variants and *cis-*only variants, an association analysis was conducted for T2D using the available phenotypic data in the UK Biobank. T2D was defined based on ICD-10 code E11 (f.41270) or self-reported diagnosis by a doctor at ≥30 years of age (f.2443 and f.2976). Individuals with type 1 diabetes ICD-10 code E10 (f.41270), self-reported doctor’s diagnosis of diabetes at <30 years-of-age (f.2443 and f.2976), gestational diabetes ICD-10 code O24 (f.41270), or self-reported doctor’s diagnosis of gestational diabetes (f.4041) were excluded from both cases and controls(DeWan et al. 2023). A total of 300,417 White British study subjects were available for analysis (N=19,964 cases and N=280,453 controls). To account for relatedness among study subjects generalized estimating equations (GEE) was applied to detect associations between each predicted NPX residual level and T2D status, adjusting for covariates age, sex, age x sex, BMI, and 20 genetic PCs.

We also analyzed 42,644 unrelated White British study subjects with measured NPX residual levels. Available for analyses were 34,650 study subjects with T2D status (N=2,556 cases and 32,094 controls). A dPWAS was conducted using logistic regression to test for an association between T2D and measured NPX residual levels, controlling for age, sex, age x sex, BMI, and 20 genetic PCs.

Following the established analytical protocol of the UK Biobank Pharma Proteomics Project(Sun et al. 2023), for association analysis between either predicted or measured protein levels and T2D, covariates were also included in the analysis to control for potential confounders which could cause spurious associations between T2D and protein expression level.

For each NPX residual level (predicted or measured), a statistical significance level of p<1.71×10⁻⁵ (a Bonferroni corrected p-value of 0.05/2,920 proteins) was used.

### Two-sample Mendelian Randomization

For those proteins which were significantly associated with T2D a two-sample MR analysis was conducted using MR-PRESSO to validate the associations between T2D and proteins. Summary statistics for T2D for ∼27 million variants were obtained from Mahajan et al., 2018(Mahajan et al. 2018) and were based on the meta-analysis of 32 T2D studies and in total contained 74,124 T2D cases and 824,006 controls of European ancestry(Mahajan et al. 2018).

For each associated NPX residual level, we obtained summary statistics for all variants that were associated (p-value<5×10^−8^) in the European discovery sample as reported by Sun et al.(Sun et al. 2023). LD clumping using PLINK2.0(Chang et al. 2015) was performed for those variants that overlapped between the significant summary statistics for each protein and the T2D summary statistics(Mahajan et al. 2018) retaining those variants within a 250 kb window of the index SNP with r²<0.1. These variants were selected as instrumental variables and used to perform MR-PRESSO(Verbanck et al. 2018). A significance level of p-value<0.05 was used to validate the associations between NPX residual levels and T2D.

## Software Availability

TransCisPredict software and documentation is available on GitHub at https://github.com/statgenetics/TransCisPredict. Additionally, all posterior weights generated from the UK Biobank are publicly available on Synapse under accession syn69052240 (https://www.synapse.org/Synapse:syn69052240/wiki/634058).

## Web Resources

Biorender: https://BioRender.com

MR-PRESSO: https://github.com/rondolab/MR-PRESSO

Pecotmr: https://github.com/StatFunGen/pecotmr

## Data Access

Individual-level proteomic, genotype array and phenotype data, which were used in this study, are available to approved researchers in the UK Biobank repository. Instructions for access to UK Biobank data are available at https://www.ukbiobank.ac.uk/enable-your-research.

## Acknowledgments

This work was support from funding from the National Institute of Health grants R01 DC017712 (S.M.L.) and A.T.D.) and R01 AG076901 (G.T.W.). Additional funding was obtained from the Urbut Family Foundation (G.T.W.). The research conducted using the UK Biobank Resource was performed under application number 32285 A.T.D.

